# Metabolic FRET sensors in intact organs: Applying spectral unmixing to acquire reliable signals

**DOI:** 10.1101/2023.05.17.541214

**Authors:** Lautaro Gándara, Lucía Durrieu, Pablo Wappner

## Abstract

In multicellular organisms, metabolic coordination across multiple tissues and cell types is essential to satisfy regionalized energetic requirements and respond coherently to changing environmental conditions. However, most metabolic assays require the destruction of the biological sample, with a concomitant loss of spatial information. Fluorescent metabolic sensors and probes are among the most user-friendly techniques for collecting metabolic information with spatial resolution. In a previous work, we have adapted to an animal system, *Drosophila melanogaster*, genetically encoded metabolic FRET-based sensors that had been previously developed in single-cell systems. These sensors provide semi-quantitative data on the stationary concentrations of key metabolites of the bioenergetic metabolism: lactate, pyruvate, and 2-oxoglutarate. The use of these sensors in intact organs required the development of an image processing method that minimizes the contribution of spatially complex autofluorescence patterns, that would obscure the FRET signals. In this article, we describe the fundamentals of intramolecular hetero-FRET, the technology on which these sensors are based. Finally, using data from the lactate sensor expressed in the larval brain as a case study, we show step by step how to process the fluorescence signal to obtain reliable FRET values.

## INTRODUCTION

The complexity of multicellular organisms relies on the division of labor among different cell types, and the resulting functional compartmentalization of specific tissues and organs (Rueffler, Hermisson, and Wagner 2012; Yanni et al. 2020). A consequence of this modular architecture of living systems is that different regions require different metabolites to either fuel biomass synthesis or satisfy specific energetic requirements. The lactate shuttle (Brooks 2018) constitutes a classic example of this metabolic compartmentalization. Besides, these requirements change over time as organisms are exposed to various environmental conditions and external stimuli. Thus, for multicellular life to be viable, metabolic pathways must be both spatially and temporally controlled (Seebacher 2018), leading to local metabolic states that might be quite different from the metabolic profiles of adjacent tissues or organs.

Most classic metabolic assays, however, lack spatial and temporal resolution, as they are based on preparing homogenates of biological material, thus preventing dynamic analysis on single samples or the direct measurement of spatial distributions of different chemical species. Therefore, despite being a basic facet of metabolic regulation, the spatial and temporal metabolic control described above is not explorable through the standard metabolic toolkit (Gándara et al. 2019). This has led to a scarcity of empirical research on the basic mechanisms that couple energy production with energy consumption in multicellular biological systems (San Martín, Sotelo-Hitschfeld, et al. 2014). Fluorescent sensors that can report metabolic states (Galaz et al. 2020) or relative abundance of metabolites in real time and in live cells (Gándara et al. 2019), on the other hand, can provide invaluable information on metabolic regulation in these multicellular systems.

Many of these metabolic fluorescent sensors are based on Förster Resonance Energy Transfer (FRET): a nonradiative transference of energy from an excited fluorophore (called donor) to another non-excited one (called acceptor) produced by dipole-dipole coupling if the fluorophores have overlapping spectra (Grecco and Verveer 2011) (Figure 1). As this phenomenon requires both fluorophores be in close proximity, with FRET efficiency decaying with the sixth power of the distance between donor and acceptor, FRET has been employed by molecular biologists as a useful trick to detect protein-protein interactions (Grecco and Verveer 2011). The most commonly employed FRET-based metabolic sensors consist in fusion proteins in which the donor and acceptor pair are linked through metabolite-specific binding domains (Figure 1 A). This configuration is known as intramolecular (donor and acceptor are different modules of the same protein) hetero-FRET (donor and acceptor are two different fluorophores) (PIETRASZEWSKA-BOGIEL and GADELLA 2011). In these sensors, the binding of a metabolite to this linker domain triggers a conformational change of the fusion protein that modifies the distance between the fluorescent cores of donor and acceptor, altering the energy transference and therefore affecting the FRET signal. Then, it is possible to associate changes in the FRET signal with changes in the relative concentrations of the monitored metabolites.

**Figure 1.**
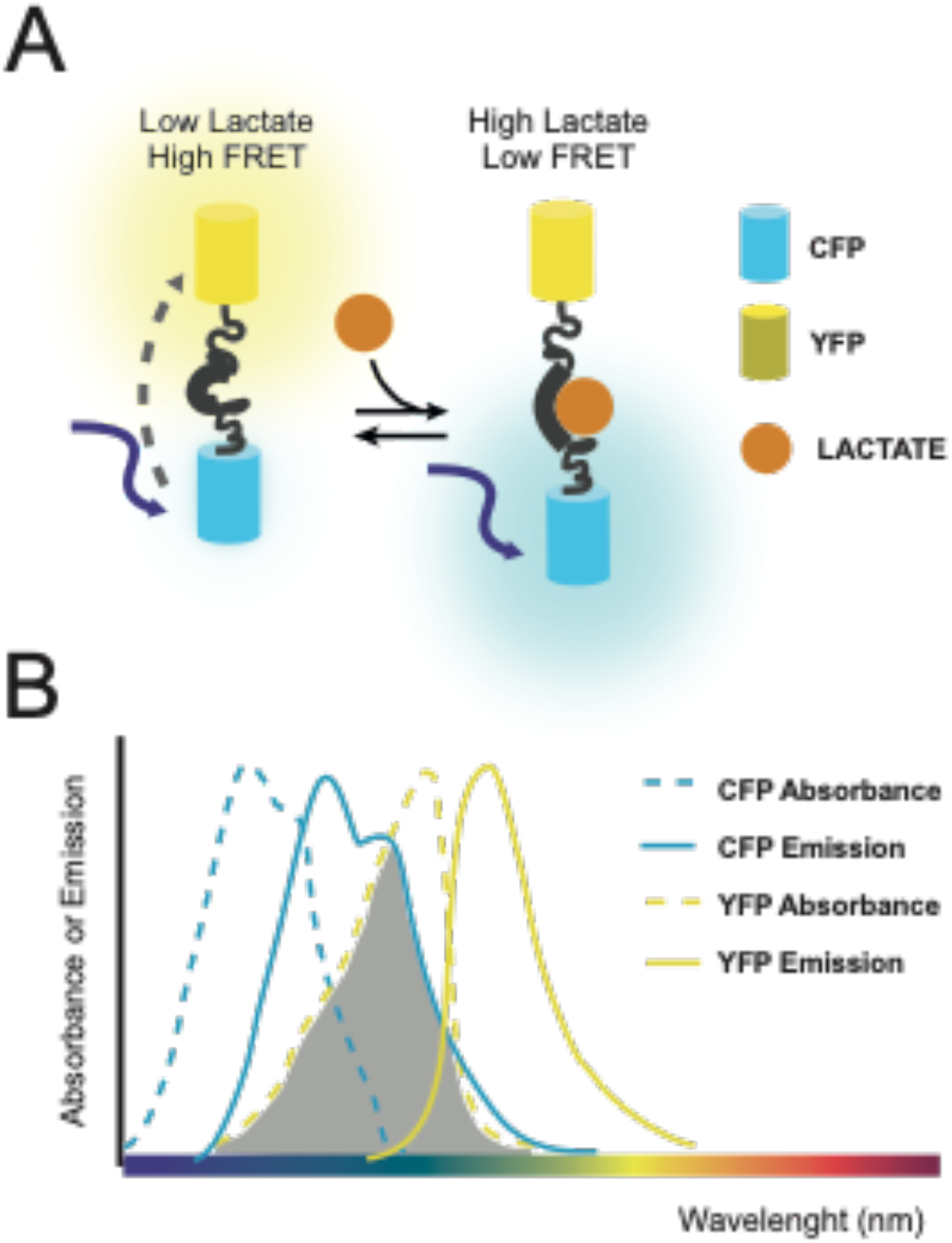
Metabolic FRET-based sensors. Illustration of the functioning of the lactate sensor Laconic (San Martín et al. 2013). A. Scheme of the molecular mechanism of the sensor. Due to the proximity of the fluorescent proteins, excitation of CFP (donor fluorophore) results in an energy tranfer to YFP (acceptor fluorophore). Union to lactate causes the fluorescent proteins to distance and thus reduces the energy transfer, and the emission of the acceptor fluorophore (YFP). B. Diagram that shows why CFP and YFP can function as a FRET pair: the emission spectrum of CFP overlaps with the absorbance spectrum of YFP (shadowed area).

Metabolic FRET-based sensors with the above-described configuration include those reporting levels of glucose (Volkenhoff et al. 2018), maltose (Fehr, Frommer, and Lalonde 2002), glutamate (Okumoto et al. 2005), NAD+/NADH (Zhao et al. 2016), lactate (Barros et al. 2013), pyruvate (San Martín, Ceballo, et al. 2014), citrate (Ewald et al. 2011), and 2-oxoglutarate (Zhang, Wei, and Ye 2013), among others. Although many of them have only been tested in cell culture or bacterial systems, during recent years these sensors have started to be used in the context of tissues (Gándara et al. 2019; Hudry et al. 2019; Morris et al. 2020).

Intact animal organs are highly complex structures, with a tightly defined tissue architecture, usually formed by several cell types, each of them fulfilling specific functions. This complexity prevents a naïve and straightforward extrapolation of imaging practices from cell cultures or bacterial suspensions to intact organs. By adapting a spectral-unmixing processing algorithm, we managed to estimate the contribution of autofluorescence to each pixel of the acquired images, and thus perform pixel-by-pixel background subtractions. This processing method removed the artefactual behavior of the FRET signals, and thus allowed us to employ these sensors for the analysis of metabolic responses in intact larval organs. In this article, we describe in detail how to obtain artifact-free FRET signals in intact organs, and we provide a step-by-step protocol for doing it.

## RESULTS AND DISCUSSION

### Imaging metabolic changes in tissues at a single-cell level

In a previous work, we have adapted the sensors Laconic (lactate), Pyronic (pyruvate) and OGSOR (2-oxoglutarate) for their use in a multicellular organism: the fruit fly *Drosophila melanogaster* (Gándara et al. 2019). To do so we cloned all three sensors downstream to UAS promoters, and then incorporated them into the fly genome through site-specific transgenesis (Bateman, Lee, and Wu 2006).

We analyzed 3rd instar larval organs (wing imaginal discs, salivary glands, fat bodies, brains and midguts) by dissection and ex-vivo imaging of the un-fixed tissues (see the protocol at Supplementary materials). We then successfully performed all the controls and analysis presented here in all three sensors (Gándara et al. 2019). In this article we will focus on the use of the metabolic sensor Laconic in *Drosophila* larval tissue. However, this protocol can be easily adapted to other species, organs, and FRET sensors.

Before measurements of the samples, we performed critical FRET controls under the selected acquisition parameters.

i. Bleed-through. Ideally, the contribution of the donor emission to the acceptor channel should be minimal. However, this cannot be assumed, but instead it must be measured. To do so, a specific sample is required in which the donor fluorophore is expressed alone. Then, we measured directly the emission of the donor in the acceptor channel, upon stimulation at the donor excitation wavelength (I^d^_so_; red variable), and found no significant bleed through (Gándara et al. 2019). However, if significant bleed-through were found, the wavelength range of the acceptor channel (acceptor maximum) should be revised. This step should be repeated for each of the different tissue types that are going to be analyzed in the experiment.
ii. Direct excitation of the acceptor fluorophore. All the measured fluorescence in the acceptor channel is assumed to come from its excitation from donor’s energy transfer. To assess experimentally whether this is the case for the specific fluorophore and microscopy conditions chosen, we imaged a sample expressing the acceptor alone (Gándara et al. 2019). As expected, we found that imaging of this sample in the acceptor channel (using the excitation wavelength of the donor) revealed no fluorescence. Finding significant emission of the acceptor by the donor excitation laser is a problem not easy to solve. Excitation with a different wavelength might be possible, but changing the fluorophore pair might be necessary.
iii. Acceptors photobleaching. In the full FRET sensor, the energy transfer itself can be verified by intentionally photobleaching the acceptor fluorophore. This is achieved by bleaching a small area with high intensity illumination, in the wavelength of acceptor excitation. The expected outcome is, upon the photobleaching, to observe an increase of the donor fluorescence (in addition to the obvious effect of a reduction of the acceptors fluorescence). To get clear results, it is convenient to perform this control under conditions where high FRET is expected.

### FRET images acquisition

Once the parameters from the technical controls have been defined for the desired tissue, images or videos of samples expressing the FRET-based metabolic sensors can be finally acquired and processed to obtain reliable FRET signals. The samples must be excited at the donor maximum absorption wavelength (Figure 1 B).

If the available confocal microscope includes a spectral module-a monochromator or similar device that allows reconstructing the whole emission spectrum of the analyzed samples, then employing this optical instrument is strongly recommended. In such case, the emission spectrum (sometimes called a lambda scan) should be acquired upon excitation with the appropriate laser wavelength for the donor fluorophore. Depending on the microscope software, it might be possible or not to define the wavelength step for the scan. In either case, after imaging the frames corresponding to the donor, acceptor and AF channels can be selected.

When using a non-spectral microscope, all samples must be imaged in three different channels simultaneously: one that includes the emission maximum of the donor, a second one, the emission maximum of the acceptor, and a third one, “AF”, at a region of the spectrum in which the fluorescence emission of the sensor is expected to be null.

### The challenge of dealing with auto-fluorescence

The calculation of a FRET signal requires values of fluorescence emission for both the donor and the acceptor fluorophores. However, measurement of the fluorescence emitted by these fluorophores is hindered by the additional contribution of spurious auto-fluorescence or background light.

Accurately estimating autofluorescence is particularly crucial in this context for two key reasons. Firstly, ratiometric calculations, such as FRET signals, are highly susceptible to the influence of autofluorescence and background noise. This happens because, mathematically, the impact of such an additional and non-relevant term in the denominator is greatly amplified. Further, given the typically small signals observed in FRET experiments, this artifact has the potential to completely erase or significantly alter the results.

Secondly, organs are complex samples that include elements -such as cells-that can have very different autofluorescence levels (Figure 2). A simple correction of autofluorescence, such as subtracting a constant value, would blur the spatial variation of the FRET signal present in these samples. Preservation of the spatial information thus require a pixel-by-pixel correction strategy.

**Figure 2.**
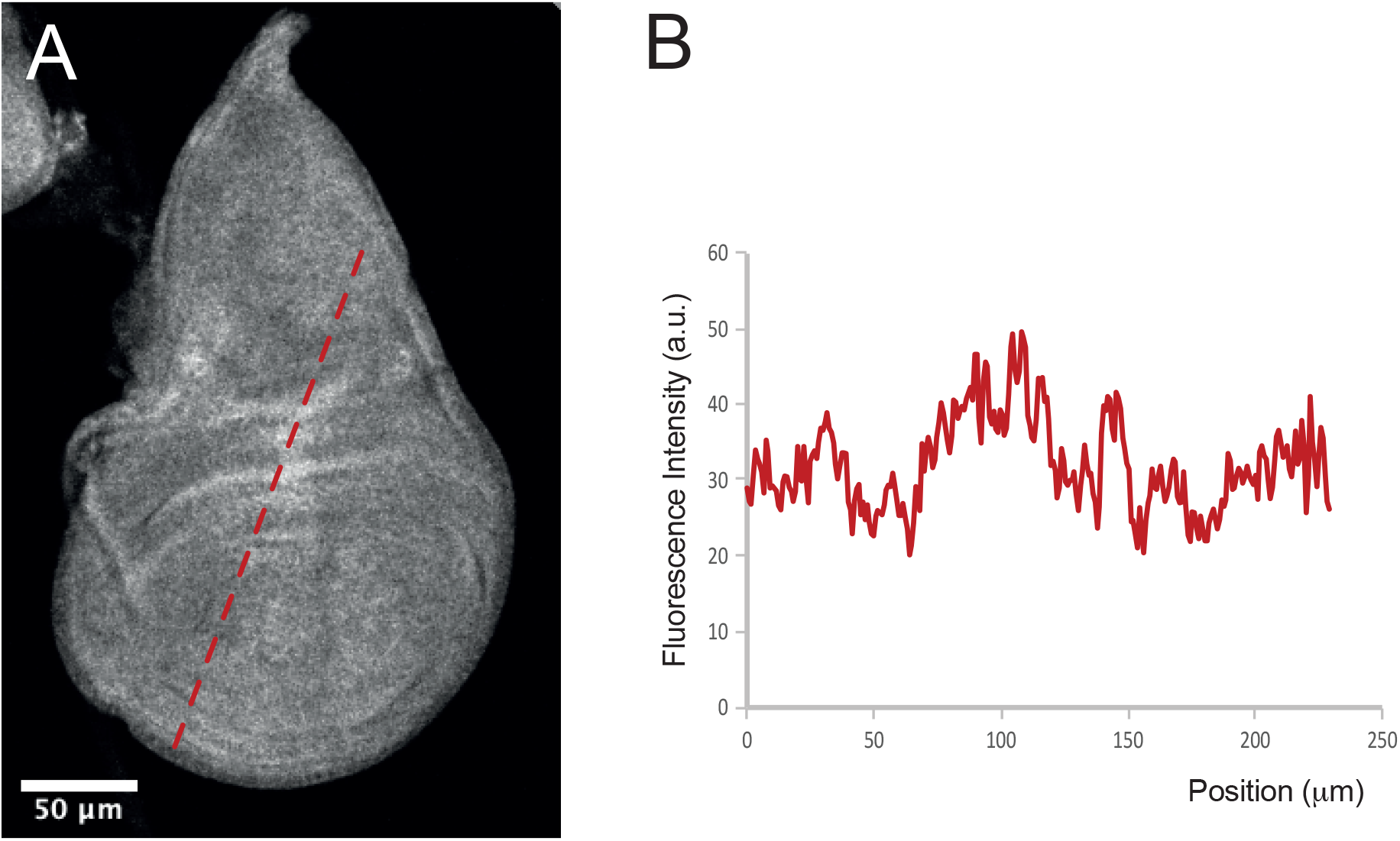
Autofluorescence varies spatially. A. Autofluorescence in a wing imaginal disc of a third instar larva. B. Fluorescence profile across the dotted red line in A.

### Spectral-unmixing for autofluorescence correction

Our first attempts to obtain a *bona fide* FRET signal when using these sensors in *Drosophila* larval organs were unsuccessful, as the FRET ratios appeared to closely reflect the expression pattern of the sensors, instead of the metabolite levels as desired (Figure 3 A-B). This effect was due to the heterogeneous autofluorescence patterns of the tissues, and the following unsuitability of commonly employed background subtraction techniques to deal with it. Thus, we implemented a linear unmixing approach for the estimation of the autofluorescence contribution to the measured fluorescence in the donor and the acceptor channels on a pixel-by-pixel basis.

**Figure 3:**
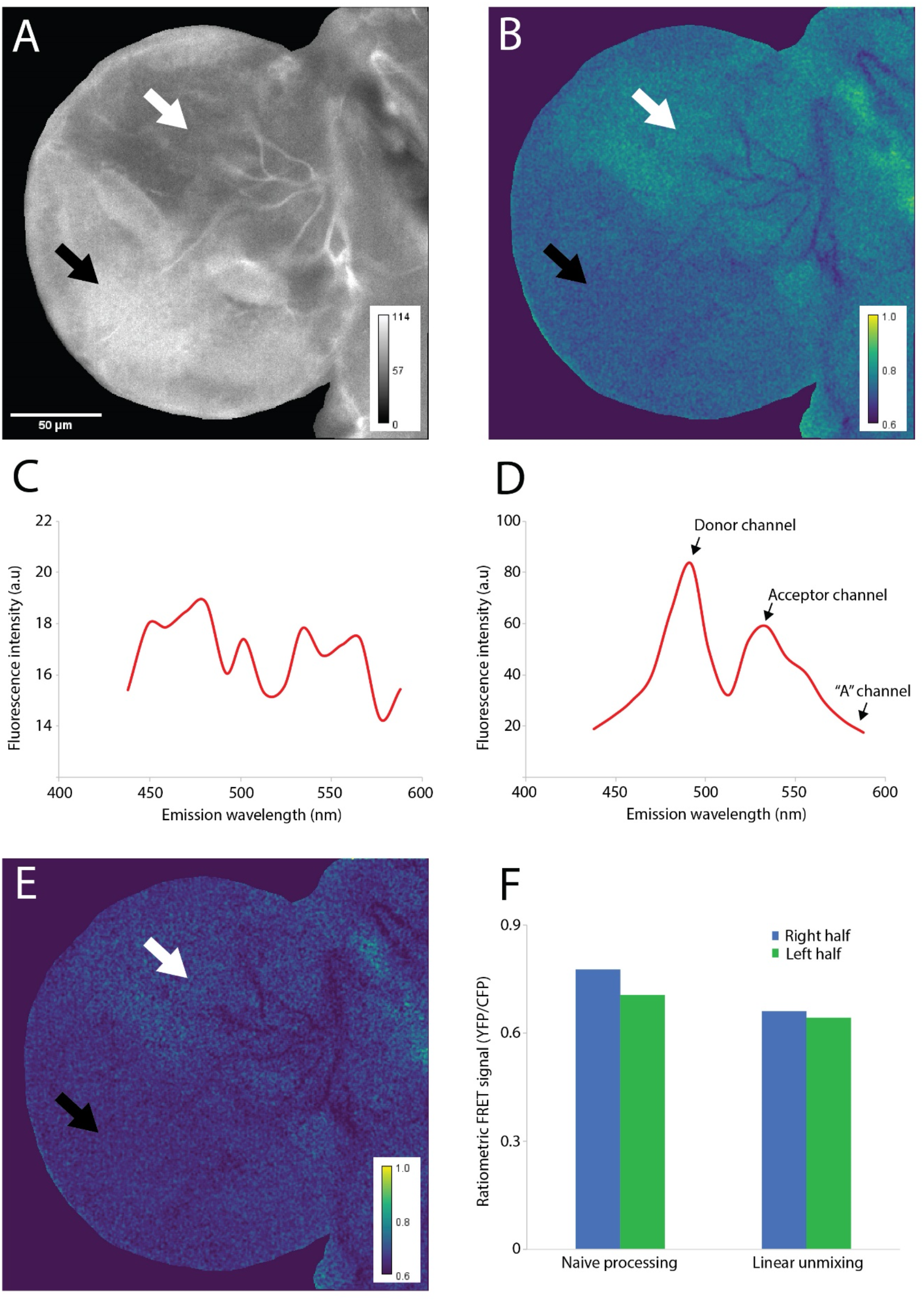
A spectral unmixing approach for correcting autofluorescence pixel-by-pixel. Larval brain lobe expressing Laconic, a lactate FRET-based sensor. A) Expression map for the lactate sensor Laconic expressed in the larval central nervous system (elav-Gal4 -> UAS-Laconic). One half of the brain lobe (black arrow) displays higher expression levels than the other half (white arrow) B) FRET map for Laconic based on its signal obtained from the image shown in (A). The straightforward ratiometric method was employed. By comparing this panel with (A), it is possible to see how the apparent FRET signal is biased by the expression pattern of the sensor. C) Autofluorescence emission spectrum of the anterior lobe of a *Drosophila* larval brain upon excitation at 458 nm (the excitation maximum of the donor fluorophore, CFP). D) Laconic emission spectrum of the anterior lobe of a *Drosophila* larval brain upon excitation at 458 nm (the excitation maximum of the donor fluorophore CFP). E) FRET map of the sensor Laconic after correcting the signal of the image shown in (A) employing the linear unmixing-based method. Note that the difference between the two halves of the lobe displaying different expression levels of the sensor (black and white arrows) was strongly reduced. F) Fret signal in the right and left halves of the lobe shown in (A), obtained either with the straightforward method used in cell culture (map shown in panel B) or after applying the linear unmixing method (map shown in panel E). The image processing algorithm reduced to a minimum the difference between the two halves of the lobe.

Independently of the method employed for image acquisition, we will refer across the manuscript to the Δλ_em_ that is expected to correspond to the maximum emission intensity window of the donor or the acceptor as the *donor channel* or *acceptor channel*, respectively.

The method assumes that the measured fluorescence emission of the donor (I_m_^d^) is composed by the fluorescence that is actually emitted by the fluorophore (I_f_^d^), plus the autofluorescence of the specific tissue in the donor channel (I_af_^d^; eq 1).

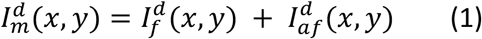

The acceptor measured emission (I_m_^a^) is similar, but has an additional component: the bleed through of the donor to the acceptor channel, i.e. the possible contribution of the donor emission due to spectral overlap (I_so_^d^; eq 2).

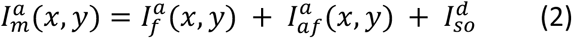

To calculate I_f_^a^ and I_f_^d^ -the actual values that are needed for obtaining the FRET signal-, the autofluorescence contribution to each channel, I_af_^a^ and I_af_^d^, must be estimated. This can be done by using a third channel in a different emission wavelength, called “AF”, where fluorescence emission from both the donor and acceptor, after stimulation at the donor excitation wavelength, is negligible (eq 3).

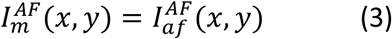

Thus, unlike I_af_^d^ or I_af_^d^, I_af_^AF^ can be measured directly (see *Image acquisition* below for the experimental instructions). As autofluorescence is characterized by a wide emission spectrum, usually reproducible for a given tissue (Desai et al. 2014), the measurement of this spectrum in tissues that do not express fluorescent sensors (Figure 1C for *Drosophila* larval brains, see *Image acquisition* below for the experimental instructions) allows the estimation of I_af_^d^ or I_af_^d^ from I_af_^AF^. Therefore, assuming that the relation between the fluorescence intensities detected in all channels is linear, it is possible to estimate I_af_^d^ or I_af_^a^ by weighting the I_af_^AF^ by a constant K (eq 4, 5).

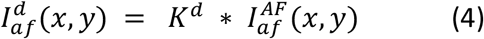

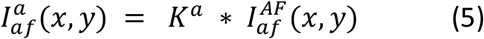

Thus, by combining eq 1 with eq 4, the *correction equation* (eq 6) for the donor channel can be obtained:

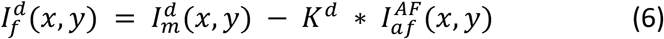

A similar combination using eq 2 and 5 will lead to the *correction equation* for the acceptor channel. Then, autofluorescence-corrected ratiometric FRET signals (F) can be calculated for each pixel of the image according to eq. 7.

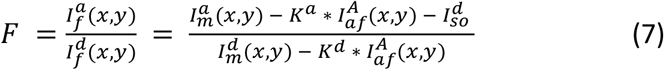

The estimation of the weighting constant K can be performed in two ways. If the experiment is design in a way that there is a region of the sample that has a similar composition to the region of interest and does not express the sensors, it can be estimated from the same image. In this case, K can be calculated as the ratio of the fluorescence in the donor or acceptor channel and the autofluorescence channel in a tissue area not expressing the sensor. This situation is ideal as the calibration is performed under the exact conditions of the experiment, and there is no need to make additional measurements. Otherwise, an independent estimation of K must be performed. This can be done by imaging organs not expressing the sensor under the same conditions than in the experiment. By using eq. 4 and 5 it should be possible to obtain values for K^a^ and K^d^ from these images.

If a spectral confocal microscope is not available (such as in Figure 3C), simple acquisition at the three-above mentioned channels will be enough (donor maximum, acceptor maximum, and “AF”).

Figure 3 shows data obtained from the anterior lobe of a *Drosophila* larval brain expressing Laconic, a lactate FRET-based sensor (Barros et al. 2013). A comparison between Figure 3 A and B shows that the image processing methods normally used in cell culture generate an artefactual FRET signal that is biased by the expression pattern of the sensor within the tissue: The two halves of the brain lobe display a clear difference of expression levels of the sensor, and the apparent (artefactual) FRET signal follows the same pattern. As shown in Figure 1F, the linear unmixing-based processing method reduces to a minimum the difference of the FRET signal between the two halves of the lobe with different expression levels of the sensor Laconic.

The image processing method described here might be more demanding than equivalent methods used to generate ratiometric FRET signals, as it requires characterizing optic properties of the tissues beforehand. In simple one-layered tissues, such as imaginal discs or the fat body, autofluorescence emission might be relatively homogeneous across space, reducing the need of estimating its contribution on a pixel-by-pixel basis. Thus, in those tissues, global background subtraction might in fact lead to a reliable FRET signal. However, Figure 3 shows that when dealing with a more complex tissue architecture, such as that of the *Drosophila* larval brain, pixel-by-pixel subtraction of autofluorescence can remove artefactual behaviors that could otherwise lead to spurious conclusions about the spatial distribution of the monitored metabolites.

A remarkable advantage of the FRET-based sensors is that their signal is reversible, which makes them suitable tools to explore dynamic facets of metabolic regulation. In this article we described how to process single images of tissues expressing FRET-based sensors. However, the same protocol can be applied to videos. By doing so, we managed to explore lactate accumulation in real time in larval tissues exposed to hypoxia *ex-vivo*, finding that the metabolic response is triggered almost immediately after lowering oxygen levels, and then, after reoxygenation, lactate levels are immediately reduced (Gándara et al. 2019).

Assessing artifact-free FRET signals that are independent of the tissue architecture becomes essential when studying the metabolic changes that underly tumor transformation, as dysregulated cell proliferation that characterizes this process can alter in a heterogeneous manner the optic properties of the organ. By using the processing method summarized here, we described in a previous work the lactogenic switch of tumors induced in wing imaginal discs. We found that the engagement on a Warburg-like metabolic switch is strongly dependent on the genetic basis of the tumor, as well as on tumor progression over time (Gándara et al. 2019). In sum, the processing method described here minimizes artefacts when obtaining FRET signals, increasing the robustness of the biological conclusions achieved when using FRET-based metabolic sensors in intact organs.

## MATERIALS AND METHODS

### Fly lines

The fly line expressing UAS-Laconic was generated in hause for a previous work (Gándara et al. 2019) by phiC31-mediated site-directed integration on the 58A landing site. The lines containing tub-Gal4 (5138), en-Gal4 (1973), ptc-Gal4 (2017) were obtained from the Bloomington Drosophila Stock Center (https://bdsc.indiana.edu/).

### Microscopy

The images were aquired using a Zeiss LSM510 Meta Confocal Microscope with monochromator. CFP was excited at 405 nm, while YFP at 458 nm. The emisión detection wondows set for the different channels were: donor (CFP) channel, 490 +/-5 nm; acceptor (YFP) channel, 530 +/-5 nm; and autofluorescence (A) channel, 600 +/-5 nm. For a detailed protocol see Suplementary Materials.

## ACKNOWLEDGMENTS

We thank the members of the Wappner lab for the discussions, with a especial thank to Camila Behrensen. We acknowladge the support of the Intitute Leloir imaging facility, and Andrés Rossi in particular. We also thank Nicholas Raun for feedback on the procedure and ImageJ macro. This work was supported by Agencia Nacional de Promoción Científica y Tecnológica (ANPCyT) grants PICT-2015-0372 and PICT-2017-1356 to P.W.

## Notes

### Competing Interest Statement

The authors have declared no competing interest.

